# Spoken language comprehension relies on context-dependent neural weighting of syntax and prosody

**DOI:** 10.64898/2026.10.05.756828

**Authors:** Cas W. Coopmans, Andrea E. Martin

**Affiliations:** New York University, New York, NY 10003, USA; Max Planck Institute for Psycholinguistics, 6525 XD Nijmegen, The Netherlands; Donders Institute for Brain, Cognition and Behaviour, Radboud University, 6525 EN Nijmegen, The Netherlands

**Keywords:** speech, structure building, prosodic boundaries, MEG, delta, temporal response functions

## Abstract

Comprehending spoken language requires extracting syntactic structure and meaning from a continuous acoustic signal. Psycholinguistic evidence points to a key role for prosody in this process. However, because interactions between prosody and syntax are typically demonstrated using carefully controlled sentence constructions, it remains unclear whether such interactions constitute a general, default property of spoken language processing. Addressing this question requires naturalistic, broad-coverage data. Here, we combined experimental psycholinguistics with naturalistic neuroscience to investigate the joint neural encoding of syntax and prosody in MEG recordings of 26 participants (19 female) listening to narrative speech. Linguistic annotations revealed that strong prosodic boundaries tend to coincide with syntactic phrase closures, and that the strength of prosodic boundaries scales with the number of phrases closed. Using time-resolved regression with temporal response functions, we modeled how these features relate to delta-band neural activity during continuous speech processing. Our results reveal a bidirectional interaction between syntax and prosody. Prosodic marking modulates the neural processing of syntax, with phrase closures being encoded more strongly when accompanied by strong prosodic boundaries. Conversely, syntactic structure shapes prosodic encoding, such that boundary strength is neurally encoded primarily at points of syntactic phrase closure. These findings demonstrate that syntax and prosody are encoded not independently, but in a context-dependent and mutually constraining manner, highlighting their joint role in guiding structure building during naturalistic spoken language comprehension.

**Significance statement:** How the brain transforms continuous sensory stimulation into structured representations is a central question in cognitive neuroscience. Spoken language comprehension provides a particularly compelling case: as the acoustic signal alone underdetermines linguistic meaning, the brain must integrate bottom-up sensory information with top-down linguistic knowledge, such as syntax. Here, we recorded brain activity during audiobook listening to examine how syntax and prosody jointly shape neural responses to narrative speech. We find evidence for a bidirectional neural interaction: prosody modulates the neural encoding of syntax, and syntactic context shapes the neural encoding of prosody. These findings suggest that the brain flexibly integrates bottom-up and top-down information depending on how informative each cue is for extracting structured meaning from naturalistic speech.

## 1. Introduction

Spoken language conveys meaning through multiple cues that must be dynamically integrated during comprehension. Bottom-up sensory signals interact with top-down linguistic knowledge to extract structured meaning from continuous speech (Martin, 2016, 2020; Poeppel et al., 2008). A central challenge for the neuroscience of language is explaining how these cues constrain one another, and whether their neural representations are fully integrated or remain partially independent. We address this challenge by investigating how syntax and prosody – two tightly linked linguistic cues – jointly shape neural processing. We use the term prosody to refer to the structured organization of spoken language and its acoustic realization in speech. Speech contains suprasegmental features that signal prosodic boundaries via pauses, pitch changes and final lengthening (Ferreira, 1993; Nespor & Vogel, 1986; Selkirk, 1984; Wightman et al., 1992). Because prosodic boundaries often align with syntactic structure, listeners can exploit them during sentence comprehension. For example, the ambiguous utterance “she likes cooking her friends and family” is parsed differently depending on prosodic phrasing. Without strong prosodic boundaries, “her friends and family” is interpreted as the object of “cooking”, whereas prosodic boundaries after “cooking” and “friends” support a list interpretation. This illustrates how alignment between prosodic and syntactic boundaries allows listeners to use prosody as a cue for syntactic parsing (Price et al., 1991; Schafer et al., 2000; Snedeker & Truswell, 2003; Steinhauer et al., 1999).

Conversely, syntactic context affects how prosodic features are processed. Listeners evaluate prosodic boundaries relative to contextual information, such as the length of surrounding constituents (Clifton et al., 2006; Deniz & Fodor, 2017). Consistent with this, the Closure Positive Shift (CPS), an ERP response to prosodic boundaries, is sensitive to constituent length: it is elicited by boundaries following long (e.g., *<u>Winnie the Pooh</u> # sold*…) but not short constituents (e.g., *<u>Pooh</u> # sold …*; Hwang & Steinhauer, 2013), even when the acoustic cues at these boundaries are identical (Holzgrefe et al., 2013).

Such evidence for syntax-prosody interactions often comes from carefully controlled experimental paradigms using structurally ambiguous sentences. Because prosodic cues in these contexts are highly diagnostic of the intended parse, the strength of the observed relationship between syntax and prosody may be exaggerated. For instance, in the example above, prosodic boundaries after “cooking” and “friends” are necessary to convey the natural interpretation. However, most utterances in everyday speech are not ambiguous in this way and will be interpreted similarly across different prosodic realizations. Moreover, in natural speech, prosodic breaks are produced for non-syntactic reasons as well (e.g., hesitations, planning), meaning that prosody is not a uniformly reliable cue to syntax (Ferreira, 2007). Listeners must therefore infer when boundaries are syntactically informative, a challenge that is attenuated in controlled paradigms. It thus remains an open question whether syntax-prosody interactions reflect a general, default property of language processing or are primarily revealed under constrained experimental conditions (Hamilton & Huth, 2020; Nastase et al., 2020).

Recent advances in encoding approaches make it possible to address this question by analyzing neural responses to ecologically realistic stimuli, in which syntax and prosody covary naturally. To date, however, most naturalistic studies examined the neural processing of syntax (Brennan et al., 2016; Coopmans et al., 2025; Giglio et al., 2024) and prosody (Bourguignon et al., 2013; Llanos et al., 2021; Teoh et al., 2019) separately. Moreover, the few studies that manipulated both variables found dissociable neural responses, in particular in the delta band (Chalas et al., 2024; Inbar et al., 2023; though see Degano et al., 2024). As a result, the neural interaction between syntax and prosody in naturalistic spoken language comprehension remains largely unexplored.

To address this gap, we investigated how syntax and prosody jointly shape neural responses to narrative speech. Using temporal response function models, we analyzed MEG recordings collected while participants listened to audiobooks. By explicitly modeling syntactic structure and prosodic boundaries, and allowing their encoding to vary with context, we tested whether prosodic cues modulate neural encoding of syntax, and whether syntactic context modulates neural encoding of prosody. Our analyses are focused on low-frequency activity in the delta band, given its relevance for processing both syntactic and prosodic structure (Coopmans & Martin, 2026).

## 2. Materials and Methods

### 2.1 Participants

Twenty-six right-handed native speakers of Dutch (19 females, 7 males, mean age = 32.7 years, age range = 19–59 years) were tested at the Donders Centre for Cognitive Neuroimaging in Nijmegen, The Netherlands. All participants reported normal hearing, had normal or corrected-to-normal vision, and had no history of language-related impairments. They provided written informed consent before the start of the experiment, which was approved by the Ethics Committee for human research Arnhem/Nijmegen (project number CMO2014/288) in accordance with the Declaration of Helsinki.

### 2.2 Materials

The narrative stimuli consisted of three fairy tales in Dutch, produced by two female speakers. The stories included a wide range of naturally occurring sentence structures with varying degrees of syntactic complexity. Across the three stories, there were 8,551 words in 791 sentences, with sentences averaging 10.8 words in length (range = 1–35, SD = 5.95). The stories were presented auditorily in 9 separate segments, ranging from 289 to 400 seconds in duration and totaling 49 minutes and 17 seconds. The loudness of each audio segment was normalized using the Ffmpeg software (EBU R128 standard). Story transcripts were automatically aligned with the recordings using the WebMAUS segmentation software (Kisler et al., 2017).

### 2.3 Analysis of the stimulus

Like in our previous work (Coopmans et al., 2025), we manually annotated the syntactic structures of all sentences in the audiobooks following an adapted version of X-bar theory, yielding binary-branching structures (Carnie, 2021). The syntactic structures contained traces of displaced elements. Displacement is the phenomenon whereby a word or phrase appears in a position different from its canonical position (e.g., the phrase in “which book did you read yesterday?”). At the latter position the displaced element is interpreted but not pronounced; it is not phonetically filled and is therefore called an empty element. Empty elements are thought to be syntactically and semantically present, but as they are not pronounced, they should not be included in an analysis of the neural encoding of prosody. Indeed, as processes of prosodic phonology appear to be insensitive to traces and other empty projections (Nespor & Vogel, 1986; Truckenbrodt, 1999), we first removed all empty elements and then collapsed all non-branching non-terminal projections, resulting in a condensed representation of syntactic structure (see Figure 1a).

**Figure 1.**
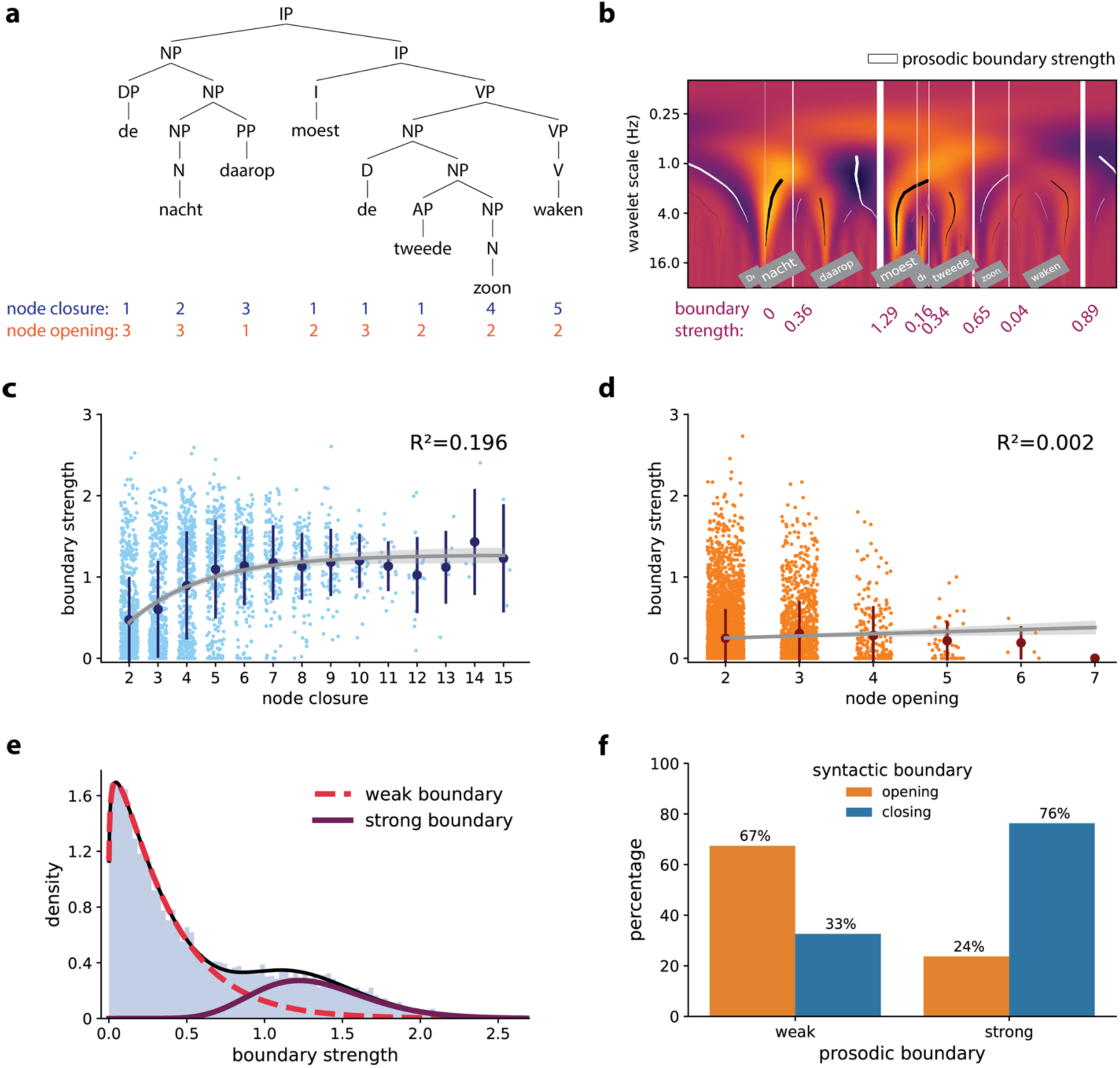
Analysis of the stimulus. **(a)** Syntactic structure for an example sentence (*De nacht daarop moest de tweede zoon waken,* lit. ‘The night thereafter must the second son watch’), with the values of node closure and node opening presented below each word. **(b)** Wavelet analysis of speech, from which prosodic boundaries are derived. The strength of the boundary is visually indicated by the thickness of the vertical white lines and is numerically presented below the wavelet scales. **(c)** Relationship between boundary strength and node closure for all phrase-final words. **(d)** Relationship between boundary strength and node opening for all phrase-initial words. The scatter points in (c) and (d) reflect individual words (slightly jittered along the x-axis for visibility), the dark circles and error bars represent mean +/− standard deviation, and the grey curve represents the fit of a regression model, with the shaded area indicating the 95% confidence interval of the fit. The R²-value indicates the proportion of variance in boundary strength explained by node closure (c) or node opening (d). **(e)** Result of the two-gamma mixture analysis, which groups all words into probability distributions of words with weak or strong prosodic boundaries. **(f)** Frequencies of phrase-initial and phrase-final words in the two prosodic clusters, derived from the ‘strong’ and ‘weak’ gamma distributions.

From the resulting tree structures, we derived two syntactic complexity metrics for each word in the audiobooks. The metric *node opening* reflects the number of phrases (or nodes in the syntactic structure) that are opened at a given word, and the metric *node closure* reflects the number of phrases that are closed (Figure 1a). As such, node closure can be interpreted as a measure of syntactic integration, whereas node opening captures the extent to which syntactic structure is incrementally constructed or anticipated. These metrics are sometimes referred to as *top-down node count* (node opening) and *bottom-up node count* (node closure), where ‘top’ and ‘bottom’ refer to the geometry of the syntactic structure (e.g., Brennan et al., 2016; Coopmans et al., 2025, 2026; Giglio et al., 2024; Slaats et al., 2024). Here, we refrain from using these terms to avoid confusion with the broader perceptual processing literature, in which *top-down* and *bottom-up* typically refer to knowledge-driven versus stimulus-driven processing, respectively.

The prosodic boundary following each word was estimated using the Wavelet Prosody Toolkit (Suni et al., 2017). This method generates metrics that have been validated against expert annotations of prosodic features (Inbar et al., 2025; Suni et al., 2017) and have been used in recent M/EEG studies testing the neural interaction between syntax and prosody (Degano et al., 2024; Inbar et al., 2023). It is an unsupervised system that estimates prosodic boundaries using a continuous wavelet analysis (CWT) on a combined signal that is composed of continuous signals representing fundamental frequency, energy, and duration. The duration signal captures both word duration and the temporal position of inter-word silent intervals (pauses), which are represented as low-valued points in the duration contour. We first estimated fundamental frequency, energy and duration, summed the time courses of these three features, and performed a CWT on the combined signal. The boundary strength value of each word is based on the output of the CWT, which is a decomposition of the signal across different scales. It is defined as the deepest trough between the strongest peaks associated with two adjacent words (Figure 1b).

A first observation is that, across the audiobooks, the number of phrases that are closed at a word is positively correlated with the prosodic boundary strength at that word. However, this relationship is not linear: prosody distinguishes words that close two through five phrases, but additional syntactic distinctions are not reflected in increasingly stronger prosodic marking (Figure 1c; see also Bannon et al., 2026). This pattern is in line with the view that prosodic structure is less deeply hierarchical than syntactic structure (Nespor and Vogel, 1986; Selkirk, 1984). To characterize this relationship, we fitted a nonlinear asymptotic model, which captured the overall trend in the data and accounted for 19.6% of the variance in boundary strength. In contrast, boundary strength is not correlated with node opening: the prosodic marking of a word is unaffected by whether it opens two, three, four or five phrases (Figure 1d). This pattern is consistent across all nine story segments (see Supplementary Figures S1 and S2), indicating that the relationship between prosodic marking and syntactic phrase closure is consistent in naturally produced speech.

To further describe the relationship between syntactic and prosodic boundaries, we conducted a mixture model analysis on the prosodic boundary strengths of all words. This analysis tests whether the observed distribution of boundary strengths can be modeled as arising from a mixture of multiple underlying distributions. Because the boundary values are strictly positive and right skewed, we modeled them using Gamma distributions (Figure 1e; see Degano et al., 2024). We compared three models: a single Gamma distribution, a mixture of two Gamma distributions, and a mixture of three Gamma distributions. Model fit was evaluated using the Bayesian Information Criterion (BIC), with lower values indicating better fit. The two-Gamma mixture model provided a substantially better fit than both the single-Gamma model (ΔBIC = 497.20) and the three-Gamma mixture model (ΔBIC = 12.55), thereby providing ‘strong’ to ‘very strong’ evidence in favor of the two-component model (Raftery, 1995). Moreover, this bimodal distribution of boundary strengths is observed in all nine story segments (see Supplementary Figure S3), confirming that it is a robust property of natural speech, as has been observed cross-linguistically (Inbar et al., 2025).

These latent prosodic distributions show a reliable association with the syntactic properties of the words. Based on the two-component gamma mixture analysis, the set of words was split into words with a *strong* prosodic boundary and words with a *weak* prosodic boundary. This split was derived from the fitted mixture model, where a boundary strength of 0.48 (corresponding to the left tail of the ‘strong’ gamma component) was used as cutoff value. Then, we fitted a logistic regression model to predict syntactic boundary type (opening vs. closing) from prosodic boundary strength. For each word, boundary type was defined according to whether more nodes were opened or closed at that word. Planned Wald tests on the model coefficients were used to assess phrase-initial (opening) versus phrase-final (closing) preferences for each prosodic set. In line with the observed positive correlation between boundary strength and node closure, words with strong boundary marking are more likely to be phrase-final words (z = –22.60, p < .001; Figure 1f). For weak prosodic marking, words were more likely to be phrase-initial than phrase-final (z = 19.03, p < .001). This correspondence suggests a natural division in prosodic boundary strength that probabilistically maps onto whether a word opens or closes a syntactic phrase (see also Degano et al., 2024).

In sum, the stimuli reveal a probabilistic relationship between syntax and prosody in naturalistic speech. Strong prosodic boundary marking tends to cue the syntactic integration demands associated with phrase-final words, whereas weak boundary marking is more characteristic of phrase openings. This relationship between syntax and prosody is probabilistic rather than deterministic because prosodic boundary presence and strength are influenced by factors beyond syntax. Non-syntactic variables such as speech rate, discourse status, and constraints on speech planning also affect prosodic structuring, resulting in considerable variability in how a given syntactic structure is prosodically realized (Bannon et al., 2026; Cole, 2015; Cutler et al., 1997; Ferreira, 2007; Shattuck-Hufnagel & Turk, 1996). Nevertheless, the observed pattern indicates that the speakers’ prosodic realizations of connected discourse systematically track fine-grained syntactic structure, rendering prosody a potentially informative cue in the speech signal (Klatt, 1975; Lea, 1972). From the listener’s perspective, prosodic structure can signal when syntactic phrases should be closed and provides information about how many closures are expected. Conversely, the syntactic parse can inform listeners when prosodic cues are likely to be informative for structure building, allowing them to selectively attend to the prosodic marking of phrase-final words. Our MEG experiment tests whether listeners capitalize on this correspondence during continuous speech processing.

### 2.4 Procedure and data acquisition

Participants were tested individually in a magnetically shielded room. They were instructed to listen attentively to the stories while remaining still and looking at a fixation cross. Following each of the nine story blocks, participants completed five multiple-choice comprehension questions with four response options each. Performance was high (mean accuracy = 87.7%, SD = 7.62%), indicating that participants attended to and comprehended the narratives.

MEG data were acquired using a 275-channel axial gradiometer CTF system with a sampling rate of 1,200 Hz. Auditory stimuli were delivered using Psychtoolbox in MATLAB (Brainard, 1997) via earphones inserted into the ear canal. Eye movements and cardiac activity were recorded using EOG and ECG electrodes, respectively. Head position was continuously tracked with three localization coils placed in custom-fitted earmolds in each ear and at the nasion (Stolk et al., 2013). During breaks between story blocks, participants were asked to adjust their head position to compensate for movement.

### 2.5 Preprocessing

The data were preprocessed using MNE Python (v0.23.1). We first downsampled the continuous MEG data to 100 Hz after applying a low-pass filter to avoid aliasing. Noisy and flat channels were automatically identified using Maxwell-based detection criteria and were subsequently interpolated. The data were then band-pass filtered between 0.5-40 Hz using a zero-phase FIR filter. Independent Component Analysis (ICA) was applied to remove artifacts resulting from eye movements (EOG) and heartbeats (ECG). For a small subset of participants, two channels failed during the recording session. These channels were interpolated to ensure a consistent number of channels across participants. The data were then segmented into epochs corresponding to the onset and offset of each story segment. For targeting slow neural dynamics, each epoch was additionally low-pass filtered at 4 Hz to extract delta-band (0.5-4 Hz) activity.

### 2.6 TRF analysis

#### 2.6.1 Stimulus features

Several base features were included in all models to account for brain responses to acoustic and word-level linguistic information. Each feature consisted of a continuous time series temporally aligned to the stimulus, with values quantifying the hypothesized cognitive response.

Acoustic processing was modeled using an eight-band gammatone spectrogram, an eight-band acoustic onset spectrogram, and a continuous measure of pitch. The gammatone spectrogram reflects moment-by-moment acoustic power, whereas the onset spectrogram captures rapid acoustic transients in the signal. Both representations covered frequencies from 20 to 5000 Hz in equivalent rectangular bandwidth space (Heeris, 2018) and were resampled to 100 Hz to match the MEG sampling rate. The onset spectrogram was derived from the gammatone spectrogram using a neurally inspired auditory edge detection algorithm (Brodbeck et al., 2020) implemented in Eelbrain (v0.37.3; Brodbeck et al., 2021).

A continuous measure of pitch was extracted using the Librosa library in Python (v0.11.0; McFee et al., 2025). We used PYIN (Mauch & Dixon, 2014), a probabilistic modification of the YIN algorithm for fundamental-frequency (F0) estimation (de Cheveigné & Kawahara, 2002). F0 was extracted at a sampling rate of 100 Hz, with the search range limited to frequencies between 80 and 500 Hz. Following previous work (Tang et al., 2017; Teoh et al., 2019), the resulting F0 trajectory was z-scored to obtain a speaker-normalized measure relative pitch.

Linguistic processing was assessed via several word-level features. The word onset feature is a binary impulse vector with value 1 at word onset and 0 elsewhere.

Word frequency was defined as the negative logarithm of word occurrence frequency in the SUBTLEX-NL database (Keuleers et al., 2010). For words absent from the database, we used the lowest observed frequency in the audiobook vocabulary.

Surprisal is the conditional probability of a word given the preceding linguistic context, quantified as the negative log of this probability (Hale, 2001). Thus, the surprisal of word *w* at position *t* is calculated via:

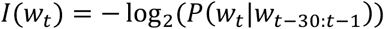

The conditional probabilities were obtained using a GPT2-Large model for Dutch from the Hugging Face library (Havinga, 2024), based on a truncated context of up to 30 tokens. GPT-2 operates on sub-word tokens, which often correspond to single words. When a word was split into multiple tokens by the GPT2 tokenizer (5.73% of all cases), we took the log probability of the first token as an index of that word’s surprisal.

Syntactic structure was captured using the features node opening and node closure, which reflect the number of syntactic constituents opened and closed at each word (Figure 1a), respectively. Prosodic structure was modeled using boundary strength, a continuous measure reflecting the strength of the prosodic boundary at each word (Figure 1b).

All word-level features were modeled on the gammatone features in terms of length and sampling rate and were time-aligned to word onsets in the auditory stimulus. Table 1 summarizes all models and included features. The acoustic spectrogram, acoustic onset spectrogram, pitch, word onset, frequency and surprisal were included in all models and are therefore collectively referred to as base features.

**Table 1.**
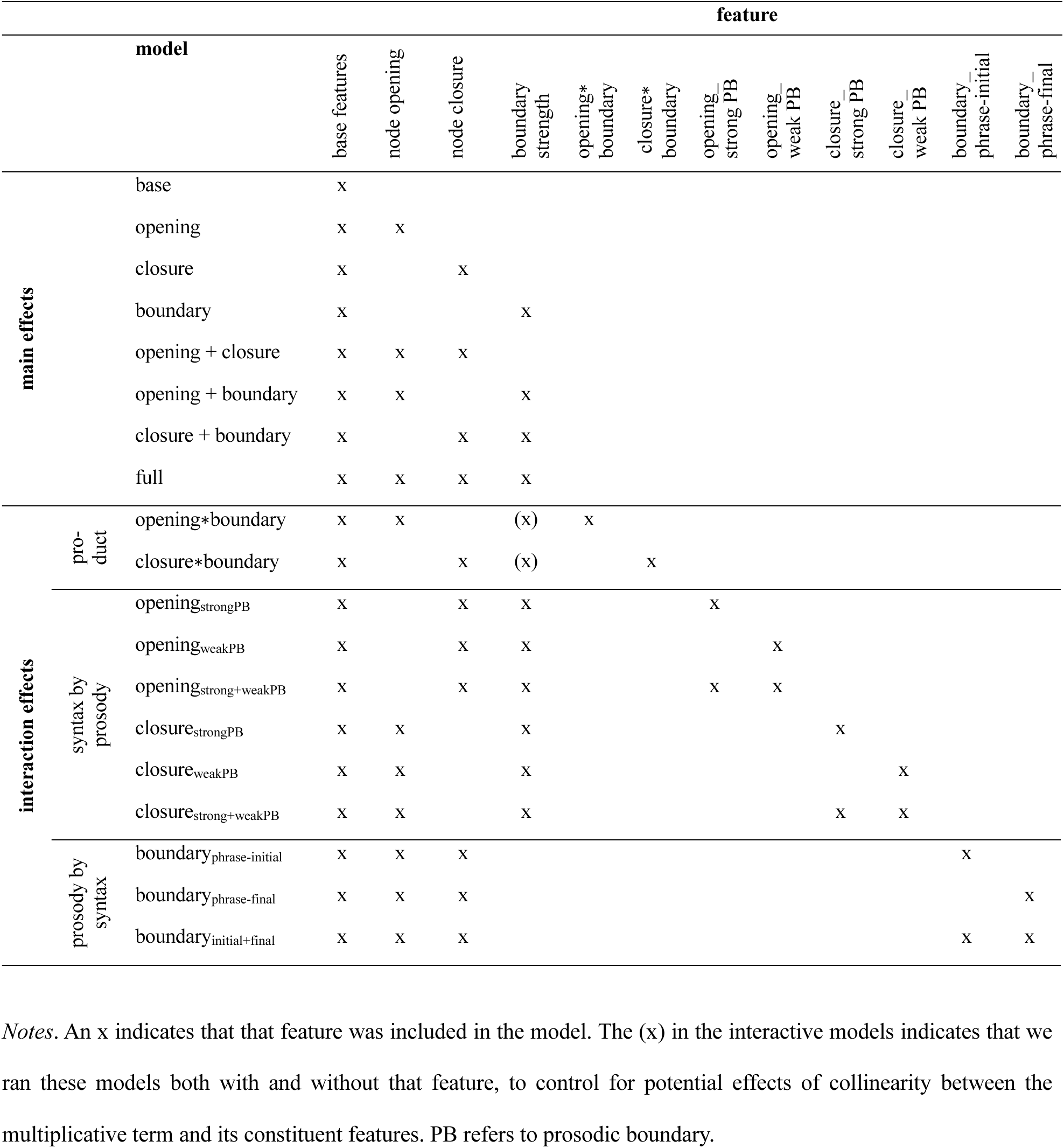
Models and included features for the main comparisons.

We reported in Section 2.3 that node closure and boundary strength are positively correlated. A high correlation between predictors in regression models can lead to multicollinearity, which refers to linear dependence among predictors and can inflate the variance of the estimated regression coefficients. This inflation is quantified by the Variance Inflation Factor (VIF), which we computed as the diagonal elements of the inverse of the Pearson correlation matrix of all word-level features (Figure 2). All VIFs were below 2, indicating no evidence of problematic multicollinearity in the present analysis (Zuur et al., 2010).

**Figure 2.**
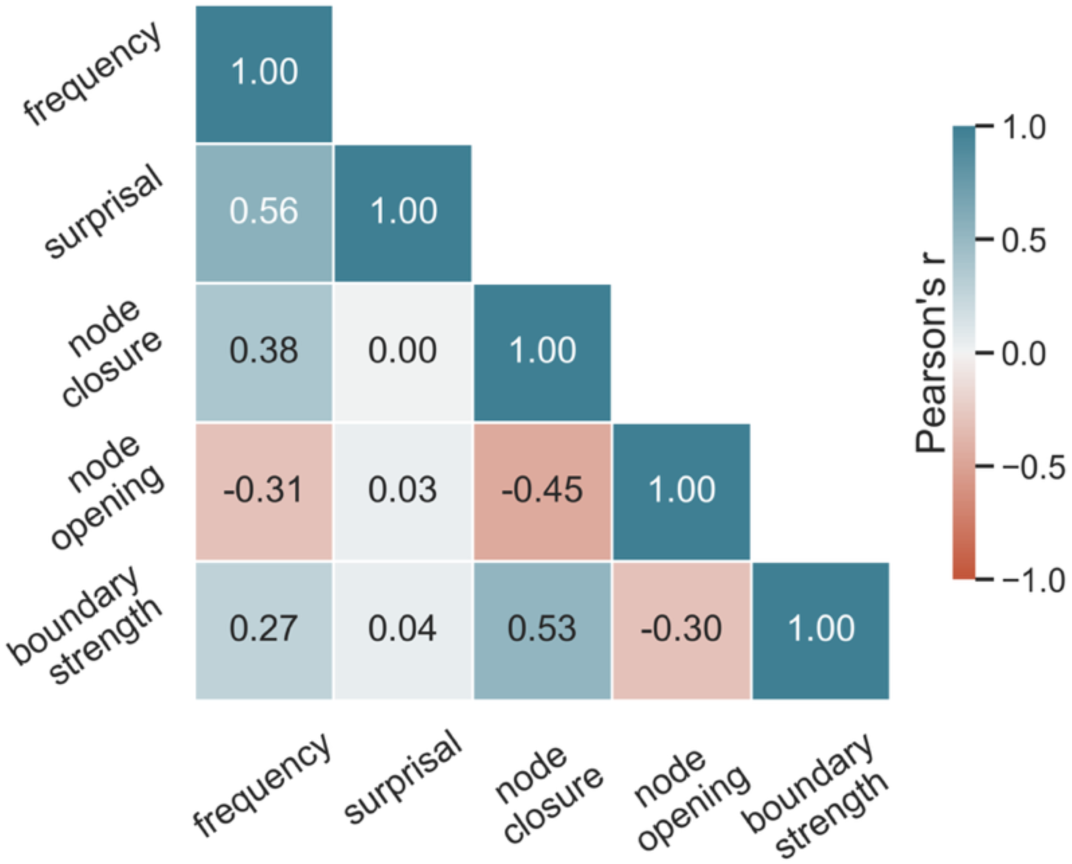
Pearson correlation of all word-level features.

#### 2.6.2 Model estimation

Temporal response functions (TRFs) were estimated for each participant and MEG sensor using Eelbrain (v0.37.3; Brodbeck et al., 2021). The MEG response at time *t*, denoted *ŷ*(*t*), was modeled as a linear combination of time-lagged stimulus features:

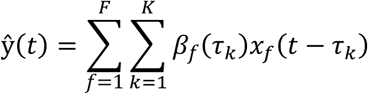

where *x_f_* denotes the *f*-th predictor time series and *β_f_*(*τ_k_*) is the coefficient of the TRF associated with predictor *f* at lag *τ_k_*. The TRF coefficient at a given latency reflects the contribution of the predictor to the MEG response at that latency.

TRFs were estimated over a temporal lag window from –100 to 1000 ms relative to stimulus onset. This window was chosen based on the latency of syntactic and prosodic responses reported in naturalistic M/EEG studies (Coopmans et al., 2025; Degano et al., 2024; Inbar et al., 2023; Slaats et al., 2024; Weissbart & Martin, 2024). To impose temporal smoothness on the estimated TRFs, we used a basis of 50-ms wide Hamming windows. Prior to model fitting, all predictors and MEG data were mean-centered and scaled by their mean absolute value to ensure comparable ranges across features.

To train and test the TRF models, we used a 5-fold cross-validation procedure. Continuous MEG recordings were concatenated across story blocks and partitioned into five equally sized folds. For each test fold, models were trained on the remaining four folds. Within the training data, an internal cross-validation procedure was used, with each of the four folds serving once as a validation set. TRFs were estimated using a boosting algorithm (Brodbeck et al., 2023; David et al., 2007) that minimizes the L1 error, with early stopping determined by validation performance. The four resulting TRFs were averaged to obtain the final model used for prediction on the held-out test fold.

This analysis yields an average TRF for each predictor in each model, as well as a measure of reconstruction accuracy for the whole model. Reconstruction accuracy refers to the fit between the predicted and the observed MEG data in the held-out test fold, quantified in terms of explained variance (R^2^). Reconstruction accuracy thus reflects the degree to which neural activity tracks variation in the stimulus features included in a given model.

#### 2.6.3 Model comparison

The main effects of the syntactic and prosodic features were evaluated by comparing the reconstruction accuracies of all TRF models that included a given feature and all models that omitted it (see Table 1). For each model, reconstruction accuracies were first averaged across sensors, yielding a single value per participant and model. These values were then analyzed using linear-mixed effects models implemented in the Statsmodels library in Python (v0.14.5; Seabold & Perktold, 2010). Each feature was represented by a dummy variable that indicated whether the feature was or was not included in the corresponding TRF model. The mixed-effects model contained the binary fixed effects node closure, node opening, and boundary strength, and included a random intercept for subject to account for between-subject variability. To avoid numerical issues due to the small magnitude of accuracy differences, the dependent variable was scaled by a factor of 10,000 prior to model fitting. This approach provides a robust estimate of each feature’s contribution to model performance that is invariant to the presence of other features.

We adopted a similar logic to evaluate the scalp distribution of the main effects. For each sensor, reconstruction accuracy was averaged across all models with a feature and compared to the average across all models without it. These contrasts were evaluated using non-parametric cluster-based permutation tests (Maris & Oostenveld, 2007) implemented in Eelbrain (v0.37.3; Brodbeck et al., 2021). For all contrasts, we applied two-tailed paired-samples t-tests at each sensor and clustered spatially adjacent sensors whose uncorrected p-value was lower than alpha = 0.05 (minimum cluster size = 5 sensors). Cluster-level statistics were computed as the sum of t-values within each cluster and compared against a reference distribution. The reference distribution was obtained from 10,000 permutations of the same data, in which the condition labels were randomly shuffled within participants. Clusters were evaluated at alpha = 0.05. Effect sizes of significant positive clusters are reported as t_clust_ (cluster-level t-value) and t_av_ (mean t-value within the cluster).

To test interactions between syntax and prosody, we first included a multiplicative interaction term, which was defined as the pointwise product between prosodic boundary strength and the two syntactic features, i.e., closure∗boundary and opening∗boundary. Within a linear TRF framework, such multiplicative predictors capture non-additive responses that scale with the joint presence of two features, such that the contribution of one feature is weighted by the instantaneous value of the other (e.g., closure weighted boundary strength or boundary-weighted node closure). Variance explained by this interaction term therefore reflects neural responses that are sensitive to the co-occurrence of syntactic structure and prosodic boundary strength, beyond the additive contributions of the individual linguistic features. We quantified the unique variance explained by the interactive feature by comparing a full TRF model including that feature to a baseline model in which that feature was omitted.

To further characterize these effects, we planned a priori to examine simple main effects of syntax as a function of boundary strength by estimating TRF models separately for words with strong versus weak prosodic boundaries (for a conceptually similar approach, see Slaats et al., 2024 and Tezcan et al., 2023). Words were split into two prosodic groups based on the gamma mixture analysis (see Section 2.3 and Figure 1e), and we tested whether the effects of the two syntactic features were different depending on prosodic boundary strength, thus aiming to capture prosody-dependent syntactic responses. Conversely, we tested whether prosodic encoding depends on syntactic context by estimating the effect of boundary strength separately for words that are phrase-initial versus phrase-final, thus capturing syntax-dependent prosodic responses.

#### 2.6.4 Analysis of the response functions

Differences between the response functions of related features were evaluated using temporal cluster-based permutation tests. The tests were conducted on the TRF coefficients averaged over sensors that showed a positive effect in the corresponding analysis of reconstruction accuracy. To compare two conditions, we applied two-tailed paired-samples t-tests at each time point. Temporally adjacent time points exceeding an uncorrected threshold of alpha = 0.05 were clustered, and cluster-level statistics were computed as the sum of t-values within each cluster. Clusters with a minimum duration of 50 ms were evaluated by comparing their cluster-level test statistic to a permutation distribution based on 10,000 random permutations of the same data. Clusters are evaluated at alpha = 0.05. To emphasize general delta-band trends in the temporal profiles, the TRFs shown in the figures were post-hoc low-pass filtered at 4 Hz. This filter is applied for visualization purposes only; all statistical analyses are performed on unfiltered TRF coefficients.

## 3 Results

We first assessed the contribution of the three main features individually. For each feature, we compared the reconstruction accuracy of all models with that feature to all models without it (see the top 8 rows in Table 1). A linear mixed-effects analysis of reconstruction accuracies averaged across sensors showed significant effects of node closure (β = 3.54e^−5^, SE = 5.84e^−6^, z = 6.11, p < .001), node opening (β = 1.11e^−4^, SE = 5.84e^−6^, z = 19.21, p < .001), and boundary strength (β = 1.46e^−5^, SE = 5.84e^−6^, z = 2.52, p = .012; Figure 3a). Given that all features improve model fit, we could evaluate their scalp topographies (Figure 3b). The effect of node closure is bilaterally distributed (t1_clust_ = 115.45, t1_av_ = 2.96, p < .001; t2_clust_ = 127.31, t2_av_ = 3.03, p < .001), as is the effect of node opening (t_clust_ = 709.98, t_av_ = 3.40, p < .001). In contrast, the effect of boundary strength is weaker and primarily distributed over right parietal-temporal sensors (t_clust_ = 24.97, t_av_ = 2.77, p = .011).

**Figure 3.**
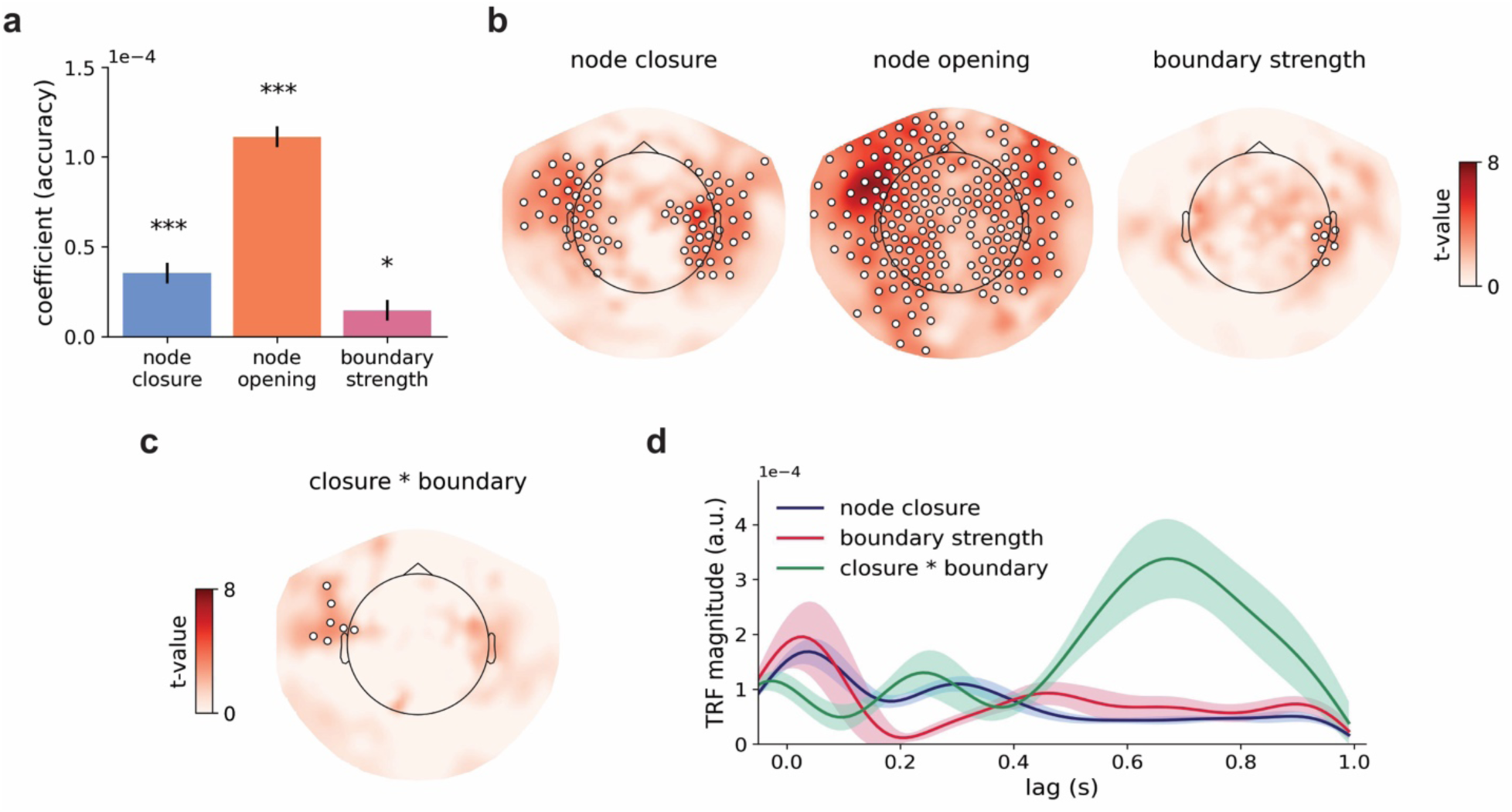
Effects of syntax, prosody, and their interaction. **(a)** Coefficients of the linear-mixed effects model of reconstruction accuracies. The coefficient of each predictor reflects its slope in a comparison between the accuracies of all models with that predictor and all models without it. **(b)** T-maps reflecting the reconstruction accuracy increases for node closure, node opening and prosodic boundary strength. The colored dots indicate sensors that were part of a significant positive cluster from the permutation t-tests. **(c)** Scalp distribution of the unique increase in reconstruction accuracy for the closure∗boundary interaction. **(d)** TRFs for node closure, boundary strength and their product, averaged over sensors that showed a significant increase in reconstruction accuracy for these features in (b) and (c).

Why is the effect of boundary strength comparatively weak? We consider two options. First, the neural responses to boundary strength might be captured by other features with which it is correlated, such as node closure or the acoustic features. Indeed, the effect of boundary strength is dependent on the presence of node closure. Adding boundary strength only improves the reconstruction accuracy of models that do not include node closure (i.e., the models base(+boundary) and opening(+boundary); cf. Table 1), which indicates that prosody and integrative syntax share variance in brain activity. Thus, the significant effects of boundary strength in Figures 3a-b are due to the fact that these comparisons include the estimate of boundary strength in models without node closure.

Another possibility is that prosody is not encoded as an independent linguistic feature but rather serves primarily as a cue to (or modulator of) other representations. Psycholinguistic evidence shows that syntactic context (e.g., constituent length) modulates whether the same acoustic cues are interpreted as a prosodic boundary or not (Clifton et al., 2006; Deniz & Fodor, 2017; Holzgrefe et al., 2013; Hwang & Steinhauer, 2013). If neural responses to prosodic boundaries are indeed context-dependent, then modeling boundary strength as a single, unitary predictor is likely to underestimate its contribution to neural activity. In the following sections, we therefore examine how neural responses to prosodic boundaries are modulated by the syntactic context in which those boundaries occur.

To explore the interactive relationship between syntax and prosody, we first tested whether including the interaction terms in the TRF models improved model fit. Here, the interaction terms are defined as the pointwise product of boundary strength and the two syntactic features, i.e., closure∗boundary and opening∗boundary. Because these interaction terms weight prosodic information by syntactic structure (and vice versa), an improvement in explained variance beyond the contribution of the individual features would indicate that syntactic and prosodic information are dynamically integrated during continuous speech processing.

Including the opening∗boundary interaction does not improve model fit. In contrast, the closure∗boundary interaction explains unique variance, even when node closure and boundary strength are already included in the base model (t_clust_ = 18.02, t_av_ = 2.57, p = .020; Figure 3c). Because multiplicative interaction terms are necessarily correlated with their constituent predictors, they can introduce multicollinearity. Indeed, variance inflation factor (VIF) analyses indicated elevated collinearity when node closure, boundary strength and their product are included in the same model (i.e., VIF > 5 for one or more predictors). Removing either node closure or boundary strength reduced VIFs below the threshold of 5 (Zuur et al., 2010). To ensure that the observed interaction effects are not driven by collinearity, we re-estimated the models after removing boundary strength. The results remain the same: the closure∗boundary interaction explained unique variance over left frontal sensors, whereas the opening∗boundary did not improve model fit.

Notably, the effect of the closure∗boundary interaction is stronger than the main effect of boundary strength alone (t_clust_ = 24.51, t_av_ = 2.72, p = .011). Moreover, its temporal response profile differs qualitatively from the response profiles of the individual predictors: whereas the TRFs for node closure and boundary strength peak early and are flat at later lags, the TRF of closure∗boundary shows a pronounced peak around 600 ms (Figure 3d). Together, these results show that jointly modeling node closure and boundary strength in a single feature captures neural variance that is not explained by either feature alone. However, the results do not yet specify the direction of the interaction: prosody may modulate the neural encoding of syntax, syntax may modulate the neural encoding of prosody, or both. In the following sections, we will address these possibilities directly.

### 3.1 Prosody modulates syntactic encoding

The prosodic modulation of syntactic encoding was assessed by splitting the set of words into words with strong and weak boundary marking and then evaluating the effects node closure and node opening for the two sets separately. As shown in Figure 4a, node closure explains variance in brain activity for words with strong prosodic marking (t_clust1_ = 26.51, t_av1_ = 2.41, p = .012; t_clust2_ = 33.99, t_av2_ = 2.61, p = .003) but not for words with weak prosodic marking. These results were further validated by a direct comparison of the effects of node closure for strongly and weakly marked words, which revealed an enhanced effect of prosodic grouping on the encoding of node closure (t = 39.89, t = 2.66, p < .001). In line with the difference in explained variance, the node closure TRF is more pronounced for words with strong boundary marking than for words with weak boundary marking (∼400–800 ms, p < .001), and it peaks late, at a lag of around 600 ms (Figure 4b). Given that the node closure response is the neural signature of syntactic integration, these results suggest that the brain is more likely to attempt syntactic integration at a word, or responds more strongly to a word’s integrative demands, when the prosodic marking of that word (probabilistically) indicates that it is a phrase-final word.

**Figure 4.**
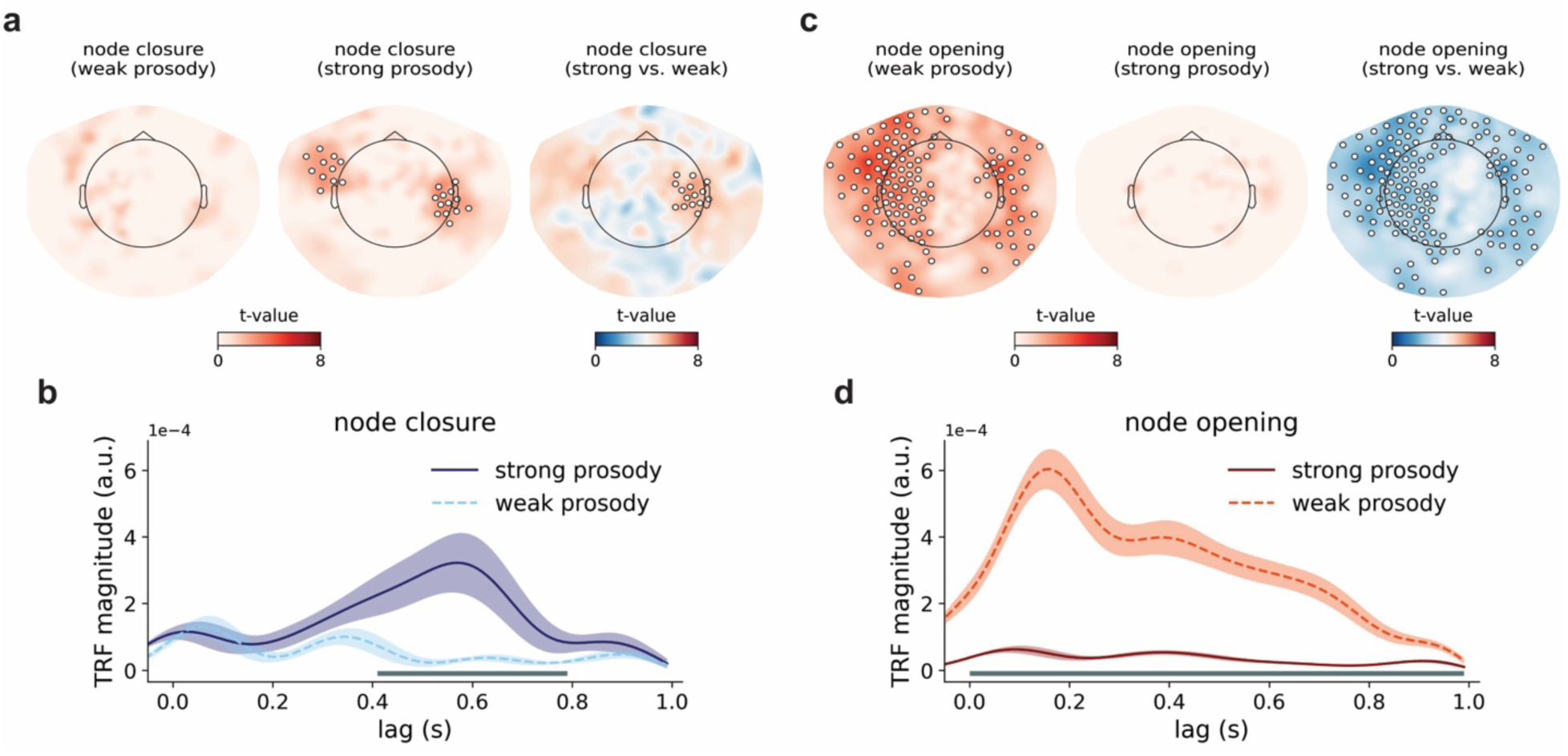
Effects of prosody on syntactic encoding. **(a)** T-maps reflecting the positive effects of node closure for words with weak and strong boundary marking, as well as their difference. **(b)** TRFs for node closure, estimated for words with strong and weak prosodic marking. **(c)** T-maps reflecting the positive effects of node opening for words with weak and strong boundary marking, as well as their difference. **(d)** TRFs for node opening, estimated for words with strong and weak prosodic marking. The response functions in (b) and (d) are averaged over the sensors that showed a significant difference in the accuracy comparisons in (a) and (c), respectively. The horizontal bars below the TRFs reflect the width of the significant clusters from temporal cluster-based permutation tests.

The results for node opening show the opposite pattern. Node opening explains neural variance for words with weak boundary marking (t_clust1_ = 287.14, t_av1_ = 2.99, p < .001; t_clust2_ = 95.98, t_av2_ = 2.82, p = .003) but not for words with strong boundary marking (Figure 4c). Here, too, the results were validated by a direct comparison of the effects of node opening for strongly and weakly marked words, which showed a stronger node opening effect for weakly marked words (t_clust_ = –464.70, t_av_ = –3.02, p < .001). The node opening TRF is more pronounced for words with weak than for words with strong boundary marking (Figure 4d). This seemingly prosodic modulation of the effect of node opening is somewhat surprising in light of the fact that the opening∗boundary interaction did not improve model fit. It might therefore reflect an effect that has a different (i.e., non-prosodic) underlying source. We will resolve this discrepancy in Section 3.3.

### 3.2 Syntax modulates prosodic encoding

Whereas the analyses in the preceding section evaluated whether the neural encoding of syntax is modulated by prosody, the current analyses assess whether the converse is true: is the neural response to prosody modulated by syntactic context? We split the set of words into phrase-initial words and phrase-final words and measured the effect of boundary strength for the two sets separately. These analyses reveal that boundary strength explains variance in brain activity for phrase-final words (t_clust1_ = 26.46, t_av1_ = 2.41, p = .007; t_clust2_ = 23.24, t_av2_ = 2.58, p = .015) but not for phrase-initial words (Figure 5a). A direct comparison of the effects of boundary strength for phrase-final and phrase-initial words showed that the difference was strongest over left frontal-temporal sensors (t_clust_ = 22.19, t_av_ = 2.47, p = .018). The differential effect of boundary strength for phrase-final and phrase-initial words shows that the effect of boundary strength is context dependent, which could explain why boundary strength (as a single, unitary predictor) does not improve model fit: there might not be a single, context-independent response to prosody in natural speech. Interestingly, the control analysis in Section 3.3.3 reveals that when the prosodic feature is split as a function of boundary strength (i.e., the effect of boundary strength for words with strong vs. weak boundary marking), this yields no improvements for either prosodic set. Thus, the result here is not merely due to be beneficial effect of splitting the prosodic feature.

**Figure 5.**
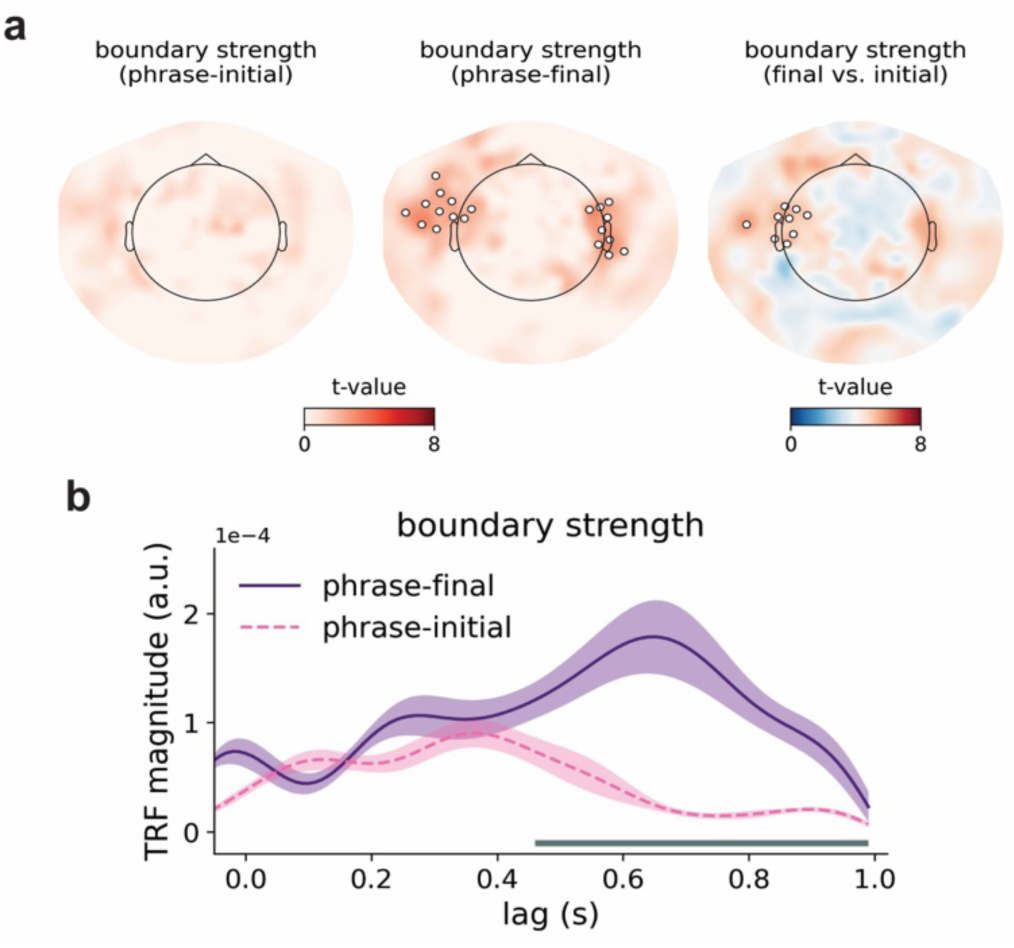
Effects of syntax on prosodic encoding. **(a)** T-maps reflecting the positive effects of boundary strength for phrase-initial and phrase-final words, as well as their difference. **(b)** TRFs for boundary strength, estimated for phrase-initial and phrase-final words. The response functions are averaged over the sensors that showed a positive difference in the comparison in (a). The horizontal bar below the TRFs reflects the width of the significant cluster from a temporal cluster-based permutation test.

Our conclusion about the context-dependent nature of prosodic processing is supported by the observation that the response functions for boundary strength are different for words in different syntactic contexts. That is, the TRFs in Figure 5b show a delayed and increased peak in response to boundary strength for phrase-final compared to phrase-initial words. This effect resembles the Closure Positive Shift (CPS) in scalp distribution, waveform shape and timing. In ERP studies, the bilateral CPS typically starts directly at a prosodic boundary (often corresponding to the offset of the pre-boundary word and the onset of the pause), has a sustained time course, and peaks roughly between 200 to 500 ms (Bögels et al., 2011; Steinhauer, 2003). The boundary-related response in Figure 5b peaks between 500 and 700 ms after word onset. Because the average duration of words in the current dataset is 240 ms, these responses correspond to an average timing of 260 to 460 ms after word offset, similar to the CPS in the ERP literature. Furthermore, recent work has linked the CPS to phase-locked delta-band activity (Henke et al., 2026; Inbar et al., 2023; Meyer et al., 2017), consistent with the delta-band time scale of the boundary response observed here.

### 3.3 Control analyses

Because of the correlation between syntax and prosody, splitting one feature based on the value of the other feature creates imbalanced sets that differ along both dimensions. For example, words with strong prosodic boundary marking are more likely to be phrase-final, whereas weakly marked words are more likely to be phrase-initial (Figure 1f). As a result, analyses that compare syntactic encoding across prosodic sets, or prosodic encoding across syntactic sets, may inadvertently confound the effect of one feature with differences in the other. Apparent effects of prosody on syntactic encoding (Figure 4), for instance, could therefore reflect differences in syntactic context rather than genuine prosodic modulation. To test whether the interaction effects reported above can be explained by such imbalances, we conducted three control analyses.

#### 3.3.1. Node closure is predictive for both phrase-initial and phrase-final words

One possible explanation of the effects in Figure 4 is that they reflect differences in syntactic context rather than prosodic modulation. Specifically, if node closure is predictive primarily for phrase-final words (with high values for node closure), then the enhanced effect of node closure for strongly marked words could arise simply because strongly marked words are often phrase-final. The same argument would hold for the enhanced effect of node opening for weakly marked words.

To test this possibility, we estimated the neural encoding of node closure and node opening separately for phrase-initial and phrase-final words. As shown in Figure 6a, node closure explains unique variance for both phrase-initial (t_clust1_ = 22.97, t_av1_ = 2.87, p = .005; t_clust2_ = 24.62, t_av2_ = 2.74, p = .004; t_clust3_ = 14.87, t_av3_ = 2.97, p = .028) and phrase-final words (t_clust1_ = 20.60, t_av1_ = 2.58, p = .013; t_clust2_ = 30.86, t_av2_ = 2.57, p = .002). The different spatial and temporal profiles of these closure effects suggest that the improvements for phrase-initial and phrase-final words may reflect partially different processes. Importantly, however, the fact that node closure is significant for phrase-initial words is inconsistent with the idea that closure is only predictive for phrase-final words. Instead, these control analyses support the interpretation that prosody modulates syntactic processing: node closure is predictive for phrase-initial words but not for weakly marked words (Figure 4a), even though weakly marked words are typically phrase-initial. This indicates that it is weak prosodic marking (rather than a phrase-initial context) that suppresses the encoding of syntactic closure.

**Figure 6.**
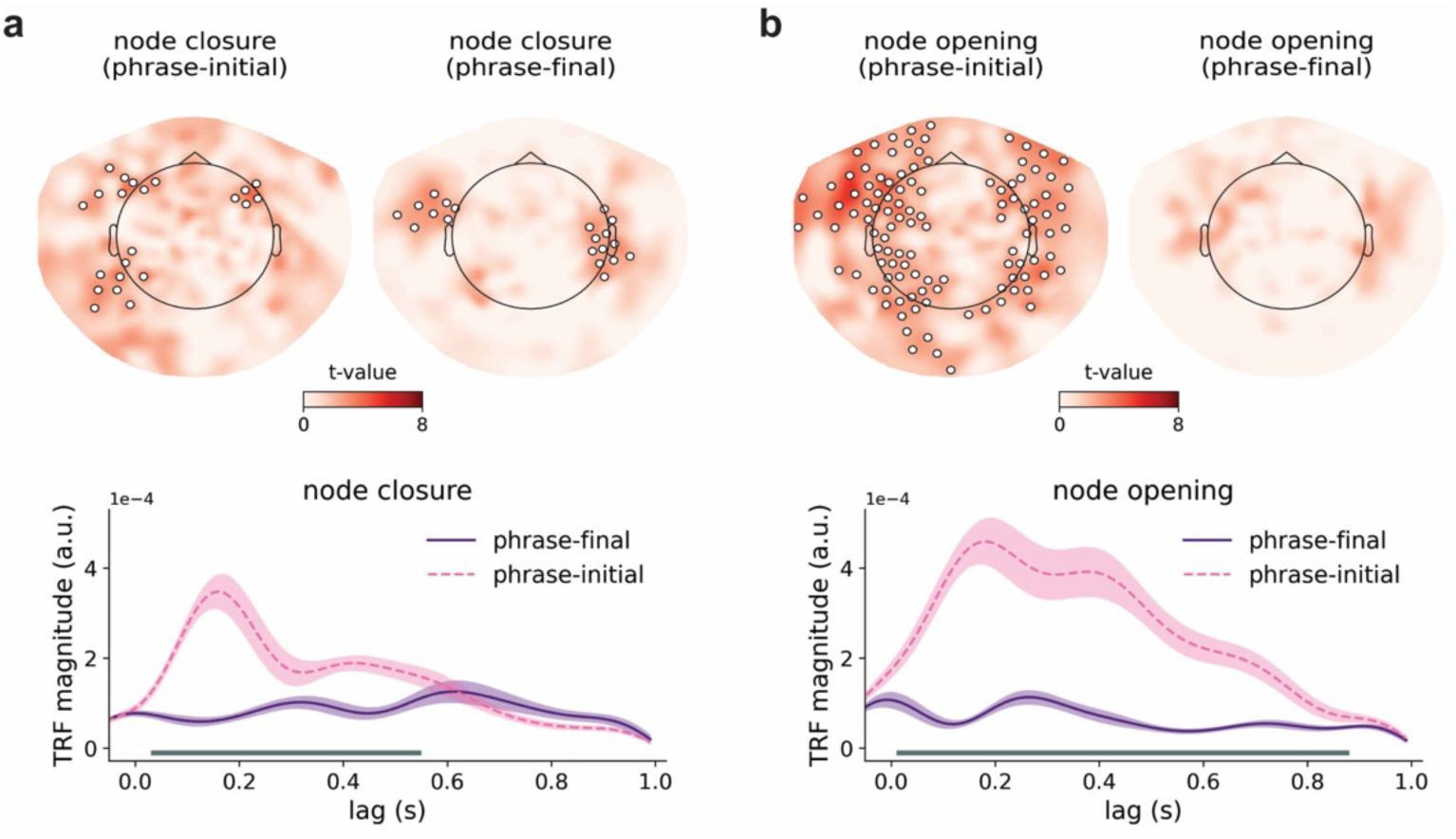
Effects of node closure **(a)** and node opening. **(b)** for phrase-initial and phrase-final words. The topographical plots reflect t-maps with improvements in explained variance, and the response functions are averaged over the sensors that showed an accuracy increase in (a) and (b), respectively. The horizontal bars below the TRFs reflect the width of the significant clusters from a temporal cluster-based permutation test.

In contrast, node opening explains variance for phrase-initial words (t_clust1_ = 205.30, t_av1_ = 2.85, p < .001; t_clust2_ = 109.02, t_av2_ = 2.73, p < .001) but not for phrase-final words (Figure 6b). This pattern mirrors the results obtained when splitting node opening by prosodic marking (Figure 4c). Moreover, the TRF of node opening for phrase-initial words resembles the TRF of node opening for weakly marked words (Figure 4d). Together, these findings indicate that node opening is primarily predictive for phrase-initial words and that its effect appears in the weak prosodic set only because weakly marked words are predominantly phrase-initial. Consistent with this interpretation, the opening∗boundary interaction did not improve model fit (Section 3), meaning that, unlike node closure, the neural response to node opening is not modulated by prosodic marking. Crucially, this pattern diverges from the syntax-prosody correspondence in the stimulus: although weak boundary marking is statistically associated with phrase openings (Figure 1f), neural encoding of node opening does not seem to track this correspondence. This shows that the brain does not merely track input statistics but projects knowledge of language onto its perceptual analysis of speech.

#### 3.3.2. Prosodic modulation of node closure persists after controlling for syntax

To directly control for the syntactic imbalance introduced by splitting words based on prosodic boundary strength, we repeated the analyses while matching the strong and weak prosodic sets for syntactic context. Specifically, we sampled from the two prosodic sets such that each set contained an equal number of phrase-initial and phrase-final words.

This procedure substantially reduced the size of the dataset, leaving 51.3% of the original words (2196 words in each prosodic set). Yet, despite this reduction in statistical power, the main result for node closure was replicated: node closure explains unique neural variance over left frontal sensors, but only for strongly marked words (t_clust_ = 20.54, t_av_ = 2.57, p = .013; Figure 7a). Although a direct comparison of the two effects showed that the effect of node closure was only marginally stronger for strongly marked words (t_clust_ = 11.13, t_av_ = 2.23, p = .076), the response functions replicate the previously observed pattern of an enhanced syntactic closure response for words with strong prosodic boundary marking (Figure 7b). Because the strong and weak prosodic sets were matched for syntactic context, this effect cannot be attributed to differences in the prevalence of phrase-final versus phrase-initial words. This control analysis therefore suggests that the effects in Figure 4a-b reflect a genuine prosodic modulation of the neural encoding of phrase closure, rather than merely stronger encoding of syntactic integration for phrase-final words.

**Figure 7.**
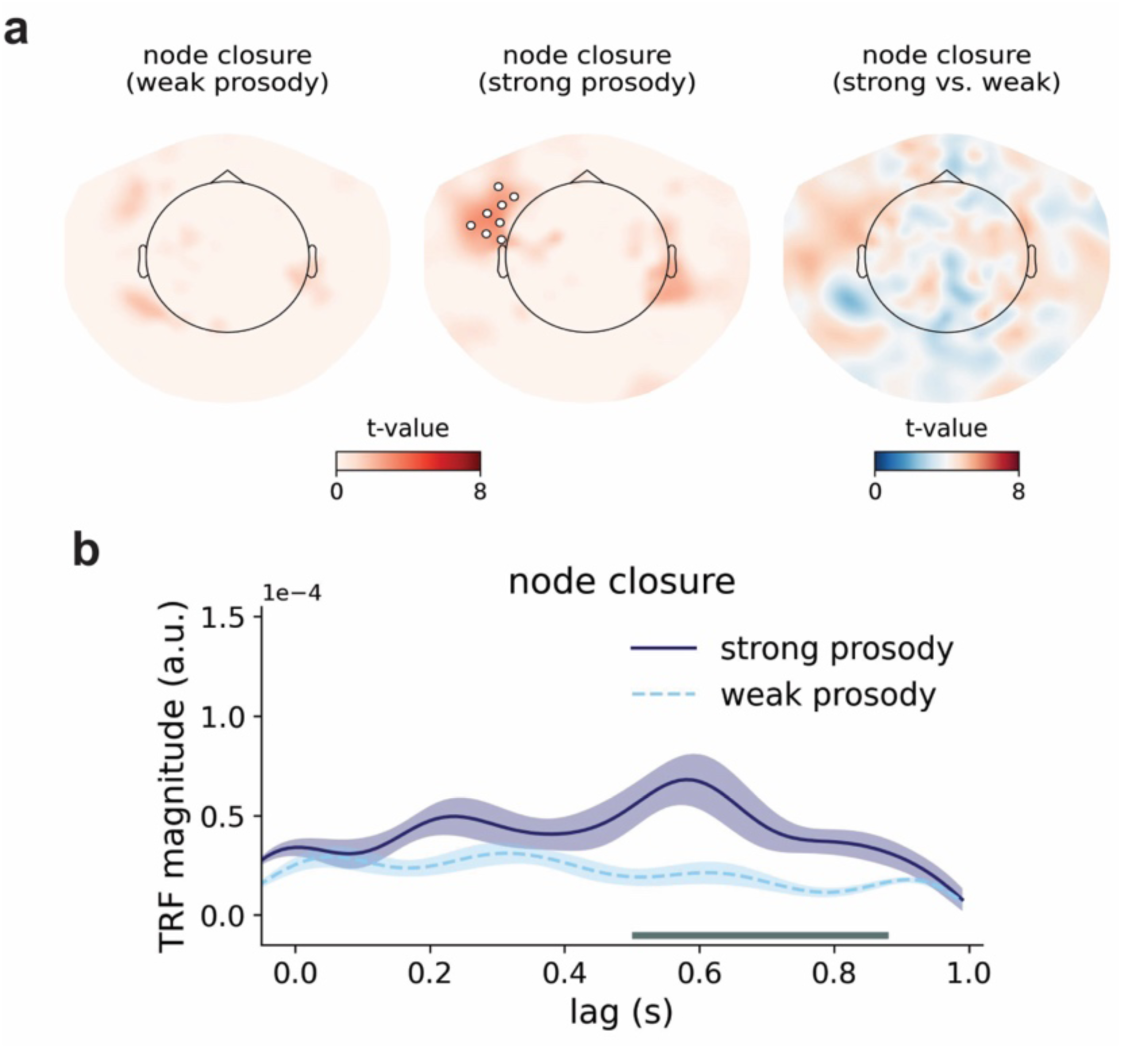
Effects of prosody on syntactic integration, while controlling for syntax. **(a)** T-maps reflecting the positive effects of node closure for words with weak and strong boundary marking, and their difference. The weak and strong prosodic sets are matched in the number of phrase-final and phrase-initial words. **(b)** TRFs for node closure, estimated for words with strong and weak boundary marking. The response functions are averaged over the sensors that showed a positive difference in the comparison in (a). The horizontal bar below the TRFs reflects the width of the significant cluster from a temporal cluster-based permutation test.

In contrast, node opening does not improve explained variance in either prosodic set. Corroborating the control analysis in Section 3.1, this suggests that the results in Figure 4c reflect the syntactic imbalance in the two prosodic sets, rather than genuine prosodic modulation: the effect of node opening was stronger for the set of weakly marked words only because the words in this set are generally phrase-initial words. Removing the imbalance by syntactically matching the prosodic sets takes away the difference in neural encoding. Together with the results of the previous control analysis as well as the non-significant opening∗boundary interaction, these findings indicate that prosodic marking selectively modulates the neural response to syntactic closure, but not the response to syntactic opening.

#### 3.3.3. Encoding of boundary strength is not enhanced by strong prosodic marking

Yet another alternative explanation follows from the relationship between syntax and prosody: phrase-final words typically have strong prosodic boundaries, whereas phrase-initial words are generally weakly marked. If neural responses to prosody are nonlinear, such that strongly marked boundaries elicit disproportionally larger responses than weak or absent boundaries (Bögels et al., 2013; Pauker et al., 2011), this could in principle account for the enhanced effects of prosody observed for phrase-final words (Figure 5), without invoking an interaction with syntax.

To test this purely prosodic account, we estimated the effect of boundary strength separately for words with strong and weak prosodic marking (for a similar conceptually approach using amplitude binning of the speech envelope, see Drennan & Lalor, 2019). If nonlinear sensitivity to prosody were driving the results, boundary strength should explain (more) neural variance in the strong set, reflecting an enhanced sensitivity to strong boundary cues. However, boundary strength does not improve reconstruction accuracy for either strongly or weakly marked words. This result indicates that merely splitting the prosodic predictor does not yield enhanced neural encoding; instead, differential effects of boundary strength emerge only when prosodic information is evaluated in relation to syntactic context (Section 3.2). Together, these results support the conclusion that the enhanced encoding of boundary strength for phrase-final words reflects an effect of syntax on prosodic encoding, rather than a nonlinear neural response to prosodic boundary strength alone.

## 4. Discussion

How the brain integrates bottom-up and top-down linguistic cues during language comprehension is a central question in the neuroscience of language. We addressed this question by investigating how syntactic structure and prosodic features interact in neural activity during audiobook listening. Analysis of narrative speech revealed a robust, probabilistic correspondence between syntax and prosody: words with strong prosodic boundaries were more likely to be phrase-closing words, and boundary strength scaled with the number of phrases closed. These statistical regularities provide perceptual cues in the acoustic-linguistic input that could be exploited by the listener’s brain during naturalistic speech processing.

We tested this using TRF models of delta-band MEG activity. Whereas recent naturalistic work has found that delta-band responses dissociate between syntactic and prosodic processing (Chalas et al., 2024; Inbar et al., 2023), our results reveal a bidirectional interaction between the two. Thus, delta-band activity does not encode syntax and prosody independently but rather integrates them in a dynamic and context-dependent manner, with each cue continuously weighted according to its relevance for extracting structured meaning from naturalistic speech.

Prior research has shown that delta-band activity is sensitive to manipulations of syntactic structure in natural speech, even when prosody is explicitly controlled (Coopmans et al., 2022, 2025; Slaats et al., 2023). This may be interpreted as suggesting that syntactic encoding is invariant to prosodic context. Here, we show that neural responses to the syntactic properties of words depend on the prosodic realization of those words. Importantly, effects of prosody on syntax were restricted to node closure, an index of syntactic integration: node closure explained more variance in neural responses to words accompanied by a strong prosodic boundary. The more anticipatory processes associated with node opening were not modulated by prosody.

Our findings are consistent with those of a recent MEG study by Degano et al. (2024). Using multivariate decoding, they showed that closing syntactic boundaries are more reliably decoded from brain responses to naturalistic speech when they are accompanied by a strong prosodic boundary. This was interpreted as evidence that prosody “boosts” the neural processing of syntax, potentially by increasing the robustness of syntactic representations. Our encoding-based approach shows that neural activity indexed by node closure – a syntactically detailed measure of closing syntactic boundaries – can be modeled accurately for strongly marked words. In contrast, when prosodic marking is weak, node closure explains less variance in neural responses, indicating that the syntactic response is either too weak or too variable to be reliably captured by this feature, hence reducing decodability.

Psycholinguistic experiments with temporarily ambiguous sentences show that comprehension is affected more by syntactically incongruent prosodic boundaries than by missing boundaries, indicating that the presence of a boundary is a stronger cue than its absence is (Bögels et al., 2013; Pauker et al., 2011). This might suggest that the absence of a boundary actively signals continuation of the current syntactic constituent (Cutler et al., 1997). However, if that were the case, weak prosodic marking should enhance the neural encoding of syntactic continuation (here indexed by node opening), which is not what we find. Instead, we suggest, in line with the correlation between boundary strength and node closure, that the brain treats the presence of a prosodic boundary as a potential cue for syntactic integration. Strong prosodic marking enables the brain to converge on the intended syntactic parse, yielding a sharpened neural representation of syntactic integration (Mai & Wang, 2023; Martin, 2020; Tezcan et al., 2026) that can be robustly estimated from neural activity (Degano et al., 2024). Weak prosodic marking instead provides a less reliable cue for parsing, which forces the listener to rely more heavily on other sources of information, such as discourse context and parsing biases. Because these can yield different outcomes in different contexts, syntactic decisions at weakly marked words will be variable. As such, prosodic marking does not merely amplify a fixed neural representation of syntax, but modulates the reliability with which syntactic structure is encoded over time.

To illustrate what this means for our encoding analysis, consider the ambiguous sentence “she saw a woman with binoculars”, which has two interpretations, each associated with a distinct prosodic contour (Nakamura et al., 2022; Snedeker & Truswell, 2003). In one interpretation – syntactically annotated as [[saw a woman][with binoculars]] – “woman” occurs at the end of the verb phrase, which is signaled by a prosodic boundary. If listeners interpret strong marking as a cue for integration, they will assign a structure in which “woman” closes multiple constituents. Neural activity associated with this interpretation is therefore well captured by the syntactic annotation given, illustrating that node closure can reliably predict neural responses in the context of prosodic marking. In an alternative analysis – represented as [saw [a woman with binoculars]] – “woman” occurs within a complex noun phrase and will typically be associated with weaker or absent prosodic marking. If weak prosodic cues are interpreted as syntactically less informative, listeners must rely on other sources of information to parse the sentence. The resulting variability in syntactic assignment reduces the consistency with which neural activity aligns with the node closure annotation, thereby lowering overall model fit for weakly marked words.

In addition, we found that the neural encoding of prosody depended on syntactic context. Boundary strength, when modeled as a unitary feature, did not reliably predict neural activity. However, splitting the feature by syntax revealed a syntactic modulation of prosodic encoding: prosodic boundaries were selectively enhanced when they coincided with syntactic phrase closure. The spatial and temporal profile of this response closely resemble that of the CPS in the ERP literature (Bögels et al., 2011; Steinhauer, 2003), which has been linked to endogenous delta-band activity (Henke et al., 2026; Inbar et al., 2023; Meyer et al., 2017). The CPS is thought to reflect prosodic structuring of the speech stream based on the integration of bottom-up acoustic and top-down linguistic cues (Bögels et al., 2011; Coopmans & Martin, 2026; Männel & Friederici, 2011), which is consistent with our finding that prosodic boundary processing is temporally aligned with syntactic integration.

It is well established that prosodic boundaries are evaluated in a context-dependent manner, rather than serving as unambiguous cue to syntactic closure or attachment. In particular, their informativeness is evaluated relative to other phonological cues, including the strength of the preceding prosodic boundaries within a global prosodic representation (Carlson et al., 2001; Frazier et al., 2006; Clifton et al., 2002). Our results extend this framework by showing that a global syntactic representation also affects the neural processing of prosodic boundaries. Given the probabilistic correspondence between syntax and prosody in naturalistic speech, this pattern suggests that the ongoing syntactic parse modulates the neural weighting of prosodic cues, enhancing prosodic processing when those cues are likely informative for structure building. In our stimuli, prosodic marking is most informative at phrase-final words (Figure 1c), consistent with enhanced prosodic encoding observed at these positions. Converging behavioral evidence shows that prosodic boundaries are perceptually more salient when they coincide with closing syntactic boundaries (Buxó-Lugo & Watson, 2016; Cole et al., 2010), consistent with the finding that prosodic processing is dynamically shaped by linguistic knowledge and experience (Gnanateja et al., 2025; Llanos et al., 2021) and by syntactic expectations in particular (Kerkhofs et al., 2007; Sammler et al., 2010).

The context-dependent encoding effects observed here have important implications for how linguistic features should be modeled in naturalistic language processing. Standard encoding approaches such as TRF models assume that the mapping between a given feature and neural activity is linear and time-invariant, such that responses can be captured by a single, unitary predictor (Brodbeck et al., 2023; Dou et al., 2024). Our results challenge this assumption by showing that the neural encoding of both syntactic and prosodic features depends on the context in which they occur. Such conditional effects are difficult to capture within a framework that assumes fixed responses independent of context (Ten Oever & Martin, 2024). Addressing this limitation will require modeling approaches that relax the typical time-invariance assumptions of TRF models, for instance by allowing response functions to vary dynamically in timing, shape and amplitude (Dou et al., 2025), or by estimating feature-specific responses separately across different contexts (Slaats et al., 2024; Tezcan et al., 2023). Further development of such interactive approaches is critical for accurately characterizing how multiple linguistic cues are simultaneously integrated during naturalistic speech processing.

In conclusion, by examining brain responses to narrative speech, we showed that listeners continuously exploit the correspondence between syntax and prosody to process both syntactic structure and prosodic features. An important extension of this work is to test these interactions in spontaneously produced speech, where prosodic breaks are distributed and realized differently than in read speech (Blaauw, 1994; Goldman-Eisler, 1972). To comprehend such everyday spoken language, the brain must not only encode prosodic cues but also determine when these sensory signals cue linguistic structure and when they do not. Explaining the flexible integration of these sources of information, especially in ecologically realistic contexts (Mai & Martin, 2025), is an important goal for the neuroscience of language.

## Acknowledgements

We thank Sophie Slaats for helpful discussions and suggestions. C.W.C. was supported by a Rubicon grant from the Netherlands Organization for Scientific Research (NWO; grant 019.241SG.007). A.E.M. was supported by a Lise Meitner Research Group “Language and Computation in Neural Systems” from the Max Planck Society, by NWO (VIDI grant 016.Vidi.188.029 and Aspasia grant 015.014.013), and by a Consolidator grant from the European Research Council (DYNALANG; ERC-2024-COG-101170162).

