## Supplementary Figures for "Spoken language comprehension relies on context-dependent neural weighting of syntax and prosody"

### Supplementary information

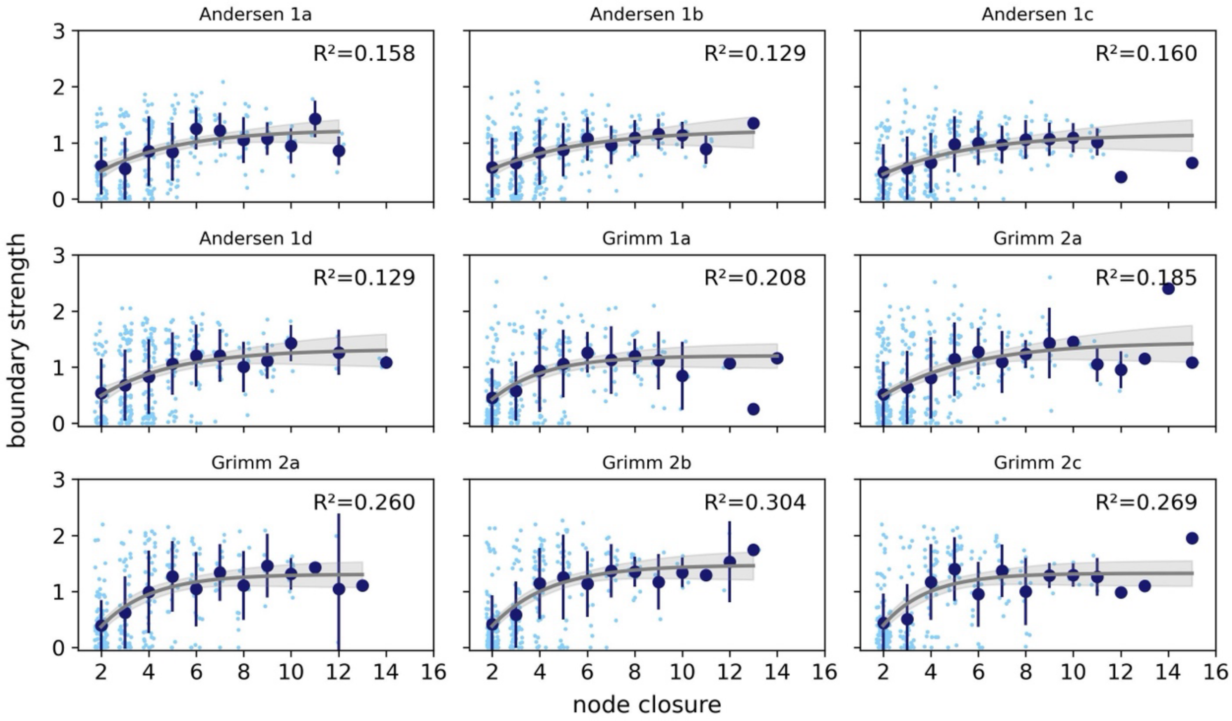


**Figure S1.** Relationship between node closure and prosodic boundary strength for phrase-final words across all nine story segments. The scatter points reflect individual words (slightly jittered along the x-axis for visibility), the circles and error bars represent mean +/- standard deviation for each value of node closure. The grey curve represents the fit of a nonlinear asymptotic model, with the shaded area indicating the 95% confidence interval of the fit. The $R^2$ value indicates the proportion of variance in boundary strength explained by node closure.

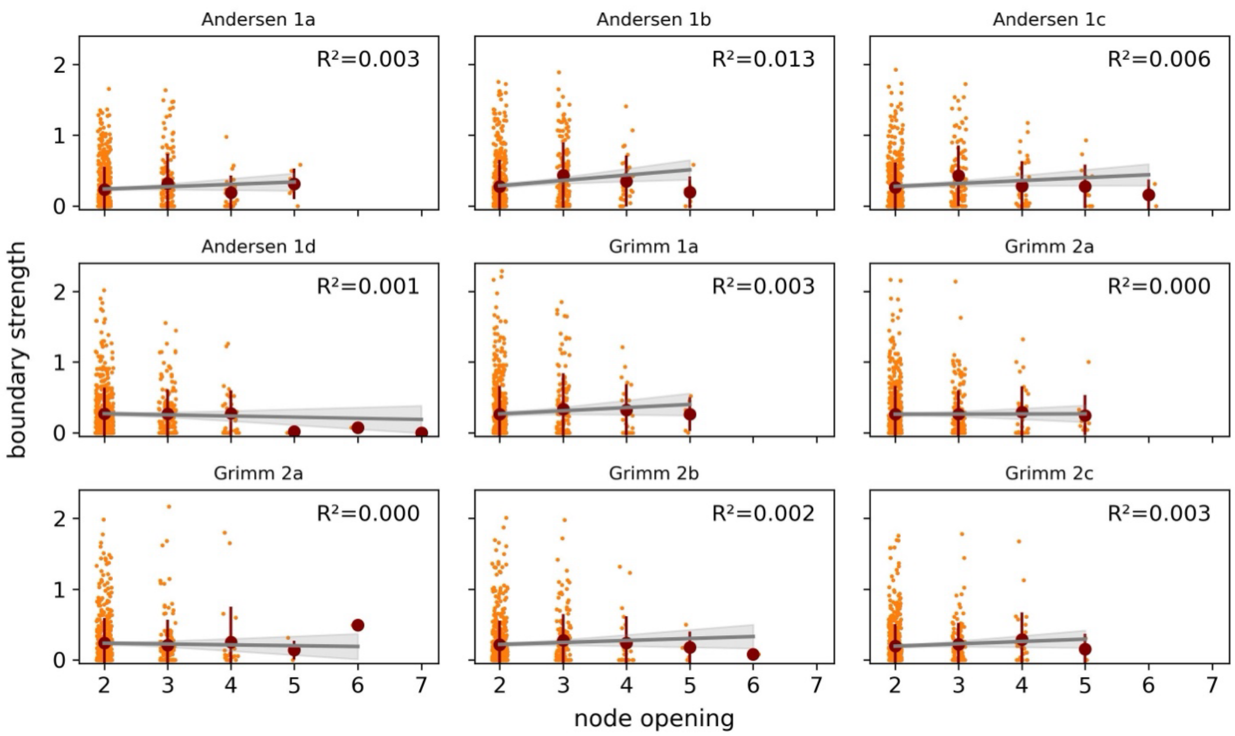


**Figure S2.** Relationship between node opening and prosodic boundary strength for phrase-initial words across all nine story segments. The scatter points reflect individual words (slightly jittered along the x-axis for visibility), the circles and error bars represent mean +/- standard deviation for each value of node opening. The grey curve represents the fit of a linear model, with the shaded area indicating the 95% confidence interval of the fit. The $R^2$ value indicates the proportion of variance in boundary strength explained by node opening.

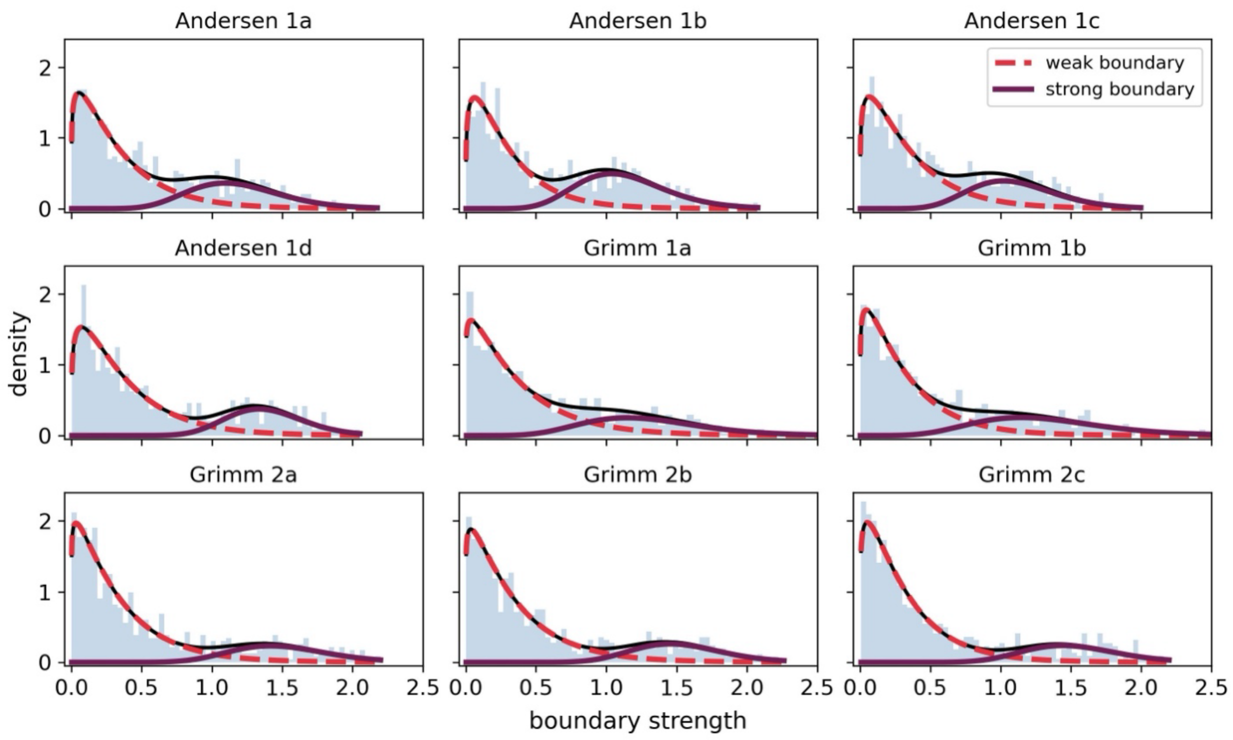


**Figure S3.** Probability distributions of prosodic boundary strengths. A mixture model analysis of each story segment reveals a bimodal distribution of boundary strengths, corresponding to words with weak and strong boundaries.
